# Brain permeable AMPK activator R481 raises glycemia by autonomic nervous system activation and amplifies the counterregulatory response to hypoglycemia in rats

**DOI:** 10.1101/749929

**Authors:** Ana M. Cruz, Yasaman Malekizadeh, Julia M. Vlachaki Walker, Paul G. Weightman Potter, Katherine Pye, Simon J. Shaw, Kate L.J. Ellacott, Craig Beall

## Abstract

AMP-activated protein kinase (AMPK) is a critical cellular and whole body energy sensor activated by energy stress, including hypoglycemia, which is frequently experienced by people with diabetes. Previous studies using direct delivery of an AMPK activator to the ventromedial hypothalamus (VMH) in rodents increased hepatic glucose production. Moreover, recurrent glucoprivation in the hypothalamus leads to blunted AMPK activation and defective hormonal responses to subsequent hypoglycemia. These data suggest that amplifying AMPK activation may prevent or reduce frequency hypoglycemia in diabetes. We used a novel brain-permeable AMPK activator, R481, which potently increased AMPK phosphorylation *in vitro*. R481 significantly increased peak glucose levels during glucose tolerance tests in rats, which were attenuated by treatment with AMPK inhibitor SBI-0206965 and completely abolished by blockade of the autonomic nervous system. This occurred without altering insulin sensitivity measured by hyperinsulinemic-euglycemic clamps. Endogenous insulin secretion was not altered by R481 treatment. During hyperinsulinemic-hypoglycemic clamp studies, R481 treatment reduced exogenous glucose requirements and amplified peak glucagon levels during hypoglycemia. These data demonstrate that peripheral administration of the brain permeable AMPK activator R481 amplifies the counterregulatory response to hypoglycemia in rats, which could have clinical relevance for prevention of hypoglycemia.

## INTRODUCTION

Achieving more time in target blood glucose (BG) range is a daily challenge for people with diabetes. This can become increasingly challenging with tightening glycemic control using insulin treatment, which increases the risk of hypoglycemia. Moreover, disease progression and frequent exposure to hypoglycemia can lead to impaired awareness of and defective counterregulatory responses (CRR) to hypoglycemia (1).

AMP-activated protein kinase (AMPK) has emerged as a central component of cellular energy sensing over the past two decades. The enzyme is a heterotrimeric complex composed of α, β and γ-subunits, with the α-subunit containing the catalytic domain (2). There are two isoforms of the α-subunit, AMPKα1 and AMPKα2, with the latter isoform having a more prominent role in glucose sensing (3-5). This enzyme plays an important role in regulating whole body energy homeostasis through its actions in the hypothalamus (6) and pancreas (7,8). Previous studies have shown that direct pharmacological activation of AMPK in the ventromedial nucleus of the hypothalamus (VMH), an important hypoglycemia-sensing region (9), increases the response to hypoglycemia in normal (10), recurrently hypoglycemic and diabetic BB rats (11) by increasing hepatic glucose production with or without concomitant increases in glucagon and epinephrine levels. Moreover, suppression of AMPK activity using shRNA diminishes the glucagon and epinephrine response to hypoglycemia (12). Recurrent glucoprivation in rats leads to diminished AMPK activation in hypothalamic nuclei during hypoglycemia (13), suggesting that at least in part, recurrent hypoglycemia (RH) may lead to defective CRR through suppression of AMPK activity in the hypothalamus. Importantly, previous studies have thus far only used direct injection of AMPK activators into the brain (10), which is not a viable therapeutic option for humans. Rigel Pharmaceuticals (CA, USA) has developed novel AMPK activating compounds with a similar mechanism of action to metformin (complex I inhibition) (14) but with greater potency). One novel compound, R481, exhibits CNS-permeability and has a positive brain:plasma distribution. We used this unique compound to test the hypothesis that peripheral delivery of a brain-permeable AMPK activator may improve the CRR to hypoglycemia.

## RESULTS AND DISCUSSION

### R481 potently activates AMPK signalling in vitro

To confirm that R481 activated AMPK in neuronal cells, we utilised the mouse hypothalamic glucose-sensing GT1-7 cells (3). Treatment for 30 minutes with increasing concentrations of R481 increased phosphorylation of AMPK at threonine 172 (a site required for full kinase activation) that was statistically significant at concentrations >10 nmol/L (Fig. 1A,B). Phosphorylation of the downstream AMPK substrate, ACC, was also significantly elevated by R481 (Fig. 1A,C). Despite AMPK activation, total intracellular ATP levels were not compromised by R481, even at concentrations up to 200 nmol/L (Fig. 1D). Using extracellular flux analysis, we confirmed mild mitochondrial inhibition, in the form of reduced oxygen consumption rate (OCR) in the mouse pancreatic αTC1-9 cell line (Supp Fig. 1). These data suggest that R481 potently and rapidly increases AMPK activity. This contrasts with metformin, which has weak brain permeability and requires transport through organic cation transporter 1 (OCT1), taking several hours to mildly inhibit mitochondria in cells (15,16).

**Figure 1.**
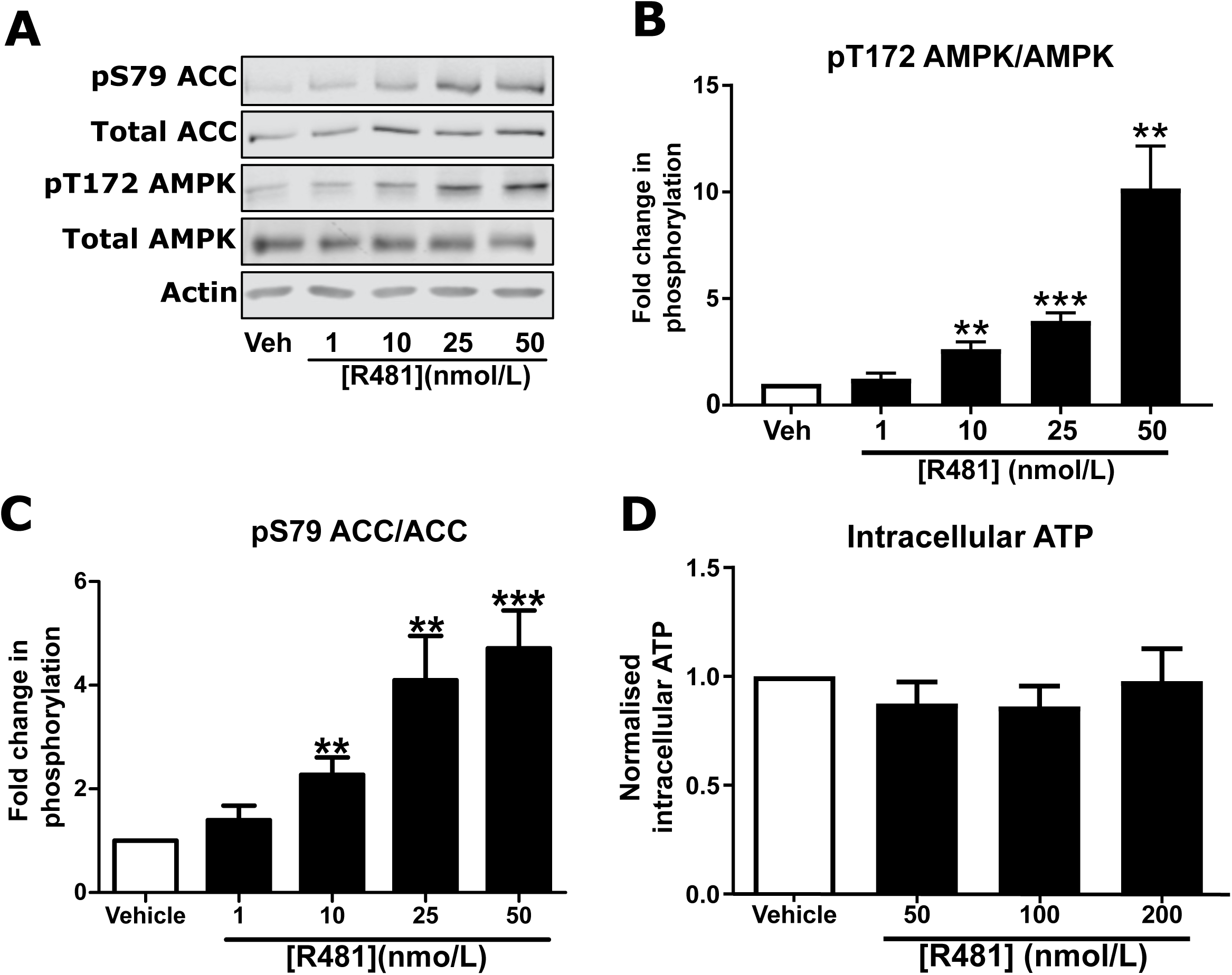
AMPK is activated in GT1-7 hypothalamic neuronal cells in response to R481. Mouse GT1-7 hypothalamic neurons treated with increasing concentrations of R481 for 30 minutes. R481 treatment resulted in a concentration-dependent increase in AMPK (pT172) and ACC (pS79) phosphorylation. **A.** Representative Western blots for AMPK (pT172), total AMPK, ACC (pS79), total ACC and Actin. Densitometric analysis of the mean pooled data for phospho-AMPK normalised to total AMPK shown in **B** (n=6) and phospho-ACC normalised to total ACC in **C** (n=8) (**P<0.01; ***P<0.001; One-sample t-test in comparison to control). **D.** Intracellular ATP levels of GT1-7 cells treated with R481, normalised to vehicle (30 mins; 0-200 nmol/L; n=6).

### R481 enters the brain but does not alter ad libitum, fasting or hypoglycemia induced feeding

Dosing studies in mice demonstrated that R481 rapidly enters the brain (Supp Fig.2A), displays a brain:plasma ratio of >3 (Supp Fig. 2B) and increases whole brain AMPK phosphorylation following bolus intravenous infusion in mice (Supp Fig. 2C). In contrast, R419 did not display significant brain permeability. As hypothalamic AMPK increases feeding (6) and leptin-induced repression of feeding requires inhibition of AMPK (17), we postulated that a brain permeable AMPK activator may increase feeding behaviour. However, R481 failed to alter *ad libitum*, fasting or hypoglycemia-induced feeding in rats (Supp Fig. 3).

**Figure 2.**
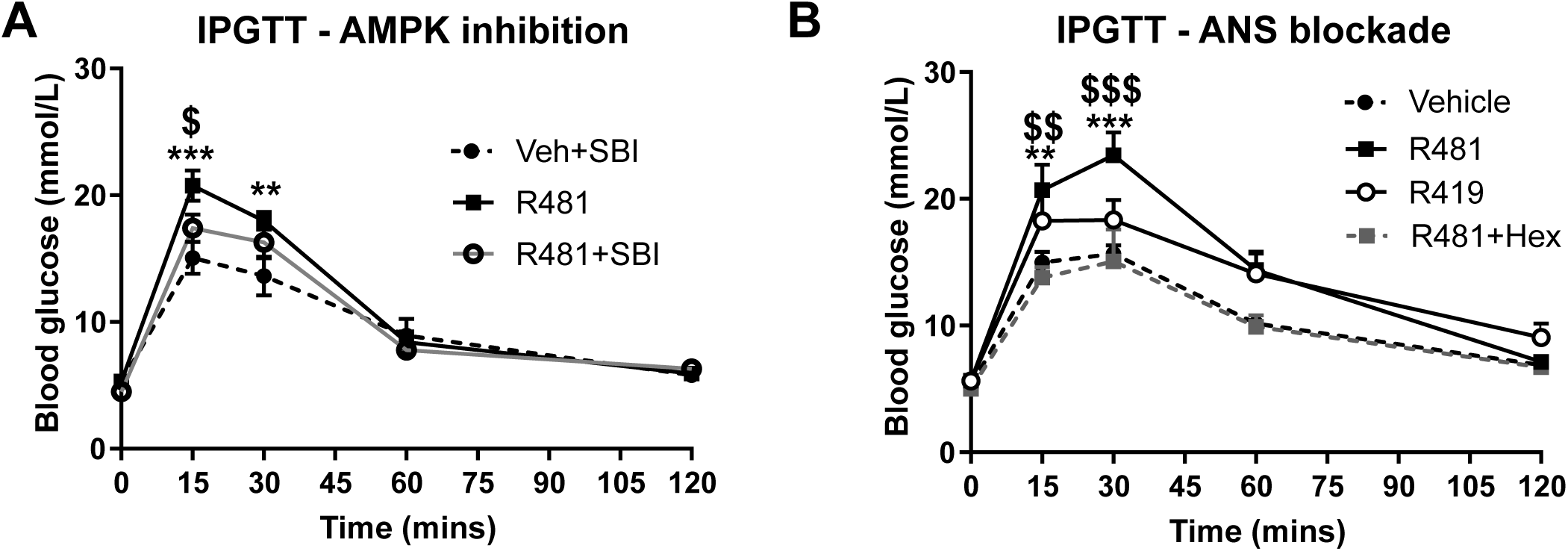
R481 induced increase in peak glycemia is attenuated by AMPK inhibitor SBI-0206965 and ANS blocker hexamethonium. Glucose tolerance tests in male Sprague-Dawley rats. **A.** After 16hr fast, rats were administered SBI-0206965 (3 mg kg^-1^; i.p.) or vehicle for 30 minutes before R481 (5 mg kg^-1^; i.p.) or vehicle (HPMC/Tween-80; i.p.) treatment together with glucose (2 g kg^-1^; i.p.). Blood glucose was measured immediately prior to R481 treatment (t=0) and after 15, 30, 60 and 120 minutes from the tail vein (SBI-0206965 n=6; R481 n=10; R481+SBI-0206965 n=8); Two-way ANOVA with repeated measures *P<0.05(drug), ***P<0.001(time), ***P<0.001(interaction) and Bonferonni’s multiple comparisons analysis, **P<0.01, ***P<0.001 for R481 against vehicle, ^$^P<0.05 for R481 versus R481+SBI-0206965. **B.** After a 16 hr fast, rats were given an glucose load (2 g kg^-1^; i.p.) alongside one of five drug treatments: vehicle (HPMC/Tween-80; n=6), vehicle with hexamethonium (50 mg kg^-1^) (Veh+Hex, n=3), R419 (20 mg kg^-1^) (n=6), R481 (20 m kg^-1^) (n=6) or R481 with hexamethonium (R481+Hex, n=4); and blood glucose measured through tail vein samples 15, 30, 60 and 120 minutes after injection. Two-way ANOVA with repeated measures, **P<0.01(drug), ***P<0.001(time), ***P<0.001(interaction), with Bonferonni’s analysis **P<0.01, ***P<0.001 for R481 against vehicle; ^$$^P<0.01, ^$$$^P<0.001 for R481 against R481+Hex.

**Figure 3.**
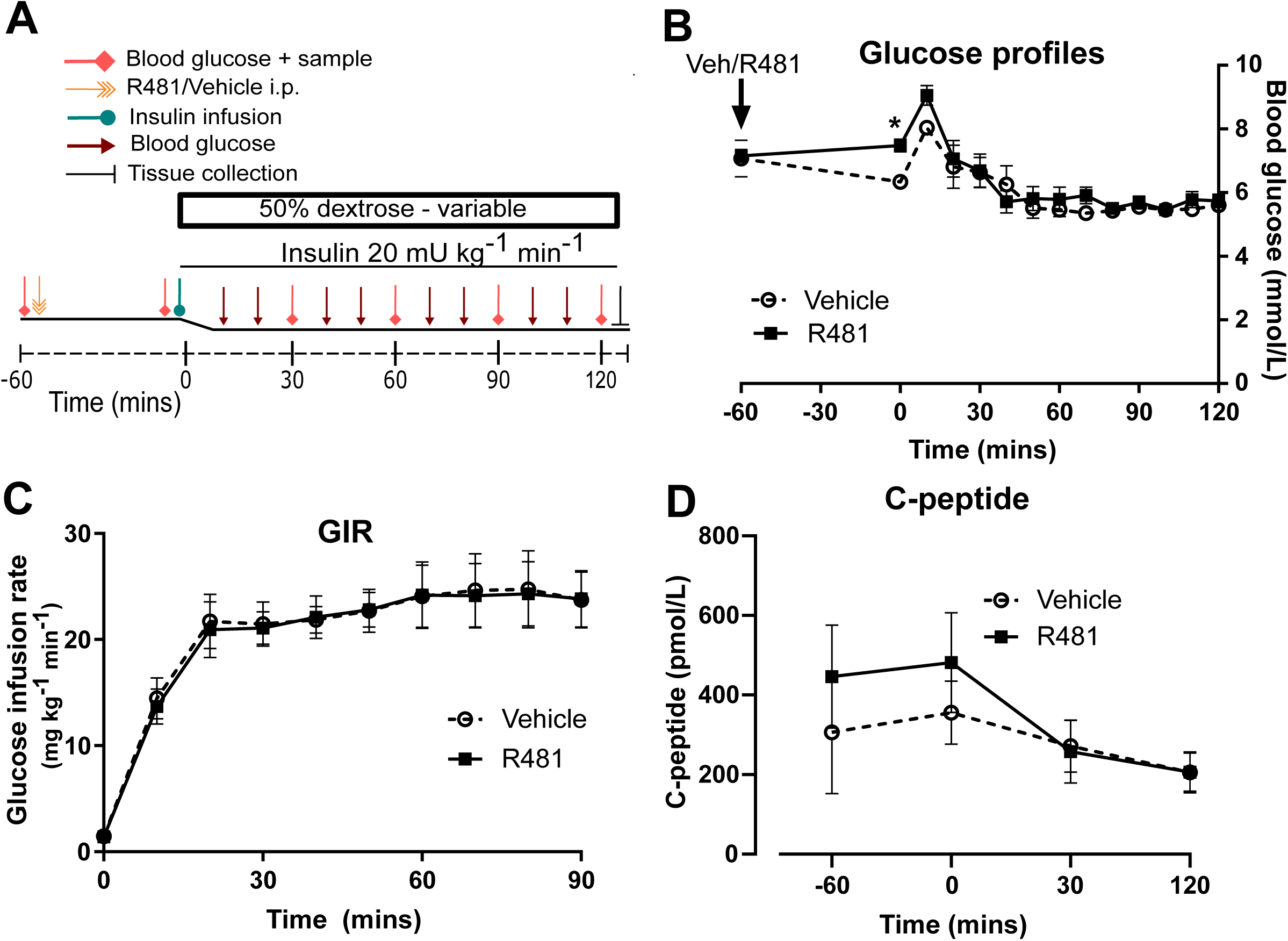
R481 does not alter glucose infusion rate during hyperinsulinemic-euglycemic clamp or alter endogenous insulin secretion. Hyperinsulinemic-euglycemic clamps performed in male Sprague-Dawley rats. **A.** Study design. **B.** Blood glucose profiles (Vehicle n=8, R481 n=8; 20 mg kg^-1^; i.p.). No overall drug effects P>0.05(drug); mixed-effects analysis of repeated measures, but *P<0.05 for R481 against vehicle at t=0 mins, using Bonferroni’s post-hoc test. **C.** Glucose infusion rate (GIR; mg kg^-1^ min-1) during the clamp using a 50% dextrose solution (Vehicle n=8; R481 n=8). **D.** Plasma C-peptide measured by ELISA (Vehicle n=8; R481 n=7).

### R481 raises peak glycemia which is attenuated by AMPK inhibitor SBI-0206965 and abolished by autonomic blockade

To determine whether R481 altered glucose tolerance, rats were treated with R481 (5-20 mg kg^-1^) simultaneously with intraperitoneal glucose. R481 significantly increased the peak glucose excursion yet glucose levels were not significantly different between groups at 2 hours, suggesting effective clearance of glucose. This transient increase in peak glucose levels was attenuated by pre-treatment with AMPK/Uncoordinated (Unc)-51-like kinase (ULK-1) inhibitor SBI-0206965 (3 mg kg^-1^; Fig. 2A), which inhibits AMPK *in vitro* at nanomolar concentrations (18). Previous studies have shown that suppression or activation of hypothalamic AMPK activity can attenuate or stimulate hepatic glucose production, respectively (19-21), indicating that hypothalamic AMPK activity regulates hepatic glucose production (HGP). Our data concur with these observations as R481 (delivered peripherally) increased glycaemia, an effect attenuated by AMPK inhibition.

The study by Kume and colleagues (22) demonstrated that activation of hypothalamic AMPK suppresses first phase glucose-stimulated insulin secretion (GSIS) through autonomic innervation of α-adrenergic pancreatic nerves. This is suggested to be a physiological response to promote glucose delivery to the brain during fasting (22), a mechanism that may also occur during hypoglycemia. Given the large and transient increase in peak glucose, coinciding with predicted first-phase insulin secretion, we pre-treated rats with pan autonomic blocker hexamethonium (50 mg kg^-1^) for 30 minutes prior to glucose tolerance testing. This drug completely abolished the effect of R481 to alter glucose tolerance (Fig. 2B). Furthermore, the R481 effect was not reproduced following treatment with non-brain permeable R419 (Fig 2B), further supporting a central action of R481 in regulating glycemia.

### R481 does not alter glucose infusion rates during euglycemic clamping

We examined whether acute R481 administration one hour prior to a hyperinsulinemic-euglycemic clamp (blood glucose target: 5.5 mmol/L) would alter glucose infusion rates. R481 (20 mg kg^-1^) or vehicle, was administered intraperitoneally 60 minutes before insulin infusion (see study design, Fig. 3A). Baseline glucose levels were slightly increased in R481 treated animals (t = −60 mins; 7.2 ± 0.2 mmol/L vs t = 0 7.5 ± 0.2 mmol/L), whereas in vehicle treated animals, BG levels decreased (t= −60 mins 7.1 ± 0.6 to t= 0; 6.3 ± 0.2 mmol/L). This produced a significant relative increase in BG at the start of the clamp in R481 treated animals (P<0.05; n=8; Fig. 3B). Glucose levels were well matched during the last 30 minutes of the clamp (Fig. 3B), with no difference in the glucose infusion rate (GIR; Fig. 3C). Contrary to our expectations, R481 had no effect on C-peptide levels (Fig. 3D) suggesting that basal endogenous insulin secretion was not altered. This suggests that the augmented peak glucose levels during the GTT are unlikely to be mediated by suppression of endogenous insulin secretion.

AMPK activators have been developed for glucose lowering in Type 2 Diabetes, largely by acting on skeletal muscle to promote glucose disposal (23,24). Moreover, the R481 analogue, R419 (non-brain permeable) activates AMPK in skeletal muscle and increases insulin sensitivity in high-fat fed mice (25). We postulated that R481 treatment may increase the GIR during the euglycemic clamp by enhancing skeletal muscle glucose uptake. However, we have no evidence of altered glucose disposal or insulin sensitivity. In our study, we examined glucose homeostasis following a single injection of the drug in lean rats, rather than chronic dosing in high fat fed mice (25), which may explain the discrepancy. However, these data suggest that the transient glucose intolerance in the GTT study was not mediated by a change to insulin sensitivity. Using glucose tracers to determine the rates of glucose appearance and disappearance will be important going forward to closely examine HGP and skeletal muscle glucose uptake.

### R481 reduces the GIR and increases glucagon levels during hypoglycemia

To expose the potential influence of central AMPK activation using R481 on CRR, we maintained rats at hypoglycaemia (2.8 mmol/L) during a 90 minute clamp study (Study design Fig. 4A). Glucose levels during the clamp were well matched between vehicle and R481-treated rats (Fig. 4B). Exogenous glucose infusion required to maintain hypoglycemia was significantly lower in R481-treated animals compared to vehicle (Fig. 4C). Plasma glucagon levels were significantly higher in the R481 treated group (Fig. 4D,E). Epinephrine levels were not modified by R481 and were undetectable 60 minutes after R481 injection (t=0; Fig. 4F,G), indicating that R481 does not alter catecholamine secretion directly. Importantly, baseline glucagon levels (during euglycemia; t=0) were not altered, suggesting that increased glucagon levels are not likely to be driving the glucose intolerance in the GTTs.

**Figure 4.**
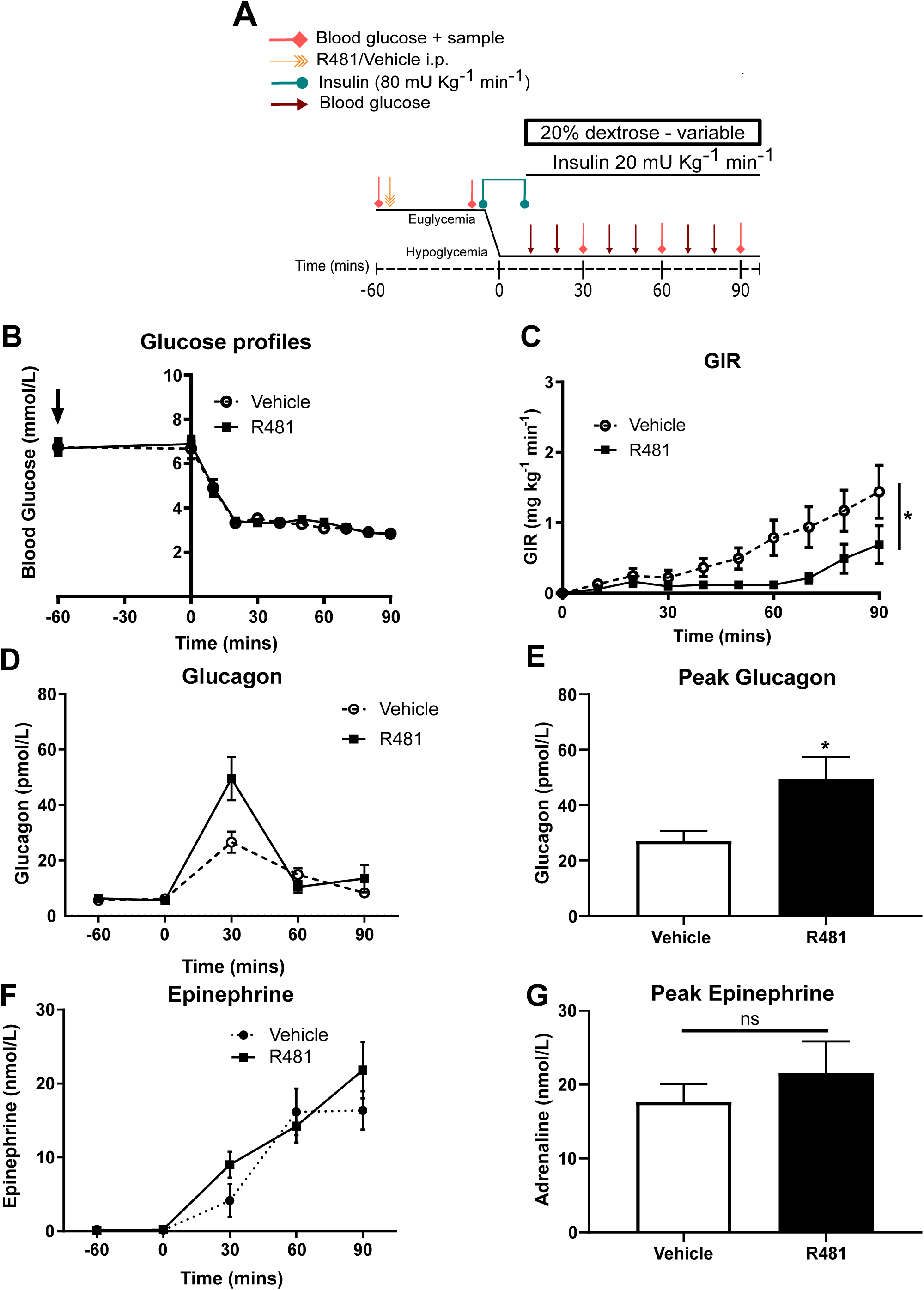
R481 delays exogenous glucose requirements during hyperinsulinemic-hypoglycemic clamp by augmenting glucagon levels during hypoglycemia. Hyperinsulinemic-hypoglycemic clamps performed in male Sprague-Dawley rats. **A.** Study design. Animals were fasted for 16 hrs. R481 (20 mg kg^-1^; i.p.) or vehicle (HPMC/Tween-80; i.p.) were administered 60 minutes before the start of the clamp. Blood glucose was measured every 10 minutes and samples collected for plasma analysis at 30 minute intervals. **B.** Blood glucose profiles before and during clamp (Vehicle n=10; R481 n=8). **C.** Glucose infusion rates (GIR; mg kg^-1^ min^-1^) during the clamp using a 20% dextrose solution. *P(drug)<0.05, ***P(time)<0.05, *P(interaction)<0.05; two-way ANOVA with repeated measures. **D.** Plasma glucagon profile with peak shown in **E**, measured by ELISA (Vehicle n=9; R491 n=9; *P<0.05, unpaired t-test). **F.** Plasma epinephrine profile with peak shown in G, meaured by ELISA (Vehicle n=8; R481 n=8; ns, not significant).

In previous studies with non-diabetic rats, direct pharmacological activation of AMPK in the VMH using AICAR, amplified HGP during hypoglycemia, without altering CRR hormones (10). In pancreatic α-cells, pharmacological and genetic activation of AMPK is sufficient to stimulate glucagon release (8). In our study, it is plausible that R481 activates an AMPK-ANS-HGP axis whilst also augmenting glucagon secretion via α-cell AMPK directly. In streptozotocin-induced diabetic rats however, VMH AICAR injection can augment both glucagon and epinephrine responses during hypoglycemia (11). Given that hyperglycemia and recurrent hypoglycemia/glucoprivation suppress hypothalamic AMPK activity (6,13) and that direct genetic suppression of VMH AMPK expression/activity suppresses glucagon and epinephrine responses to hypoglycemia (12) it is plausible that hypothalamic AMPK activity is blunted in diabetes, leading, at least in part, to defective CRR.

We demonstrate that R481 raised peak glucose levels during GTTs, without negatively impacting glucose clearance in a manner that was attenuated by AMPK inhibition and completely abolished by autonomic blockade, suggesting R481 acts centrally. Moreover, R481 did not alter C-peptide levels, suggesting that R481 does not suppress insulin secretion and likely increases glycemia by stimulating HGP, as has been previously reported following viral and pharmacological manipulation of hypothalamic AMPK activity (19-21). We also show that during hypoglycemia glucagon levels were amplified by R481. This effect could be mediated by direct activation of AMPK within the α-cell as pharmacological and genetic activation of α-cell AMPK stimulates glucagon secretion (7,8). Importantly, this indicates that the likely net effect of AMPK activation throughout the whole body is to increase glucose delivery to the brain, as previously suggested (22), indicating that at the level of the whole organism, central AMPK activation may supersede peripheral activation in a hierarchical manner, akin to that suggested for subcellular pools of AMPK (26).

In summary, our data indicate that peripheral delivery of a brain permeable AMPK activator raises glycemia, likely to protect brain function by providing more substrate for brain cell metabolism. We provide proof-of-concept that pharmacological activation of AMPK may be a suitable therapeutic target for amplifying the defence against hypoglycemia. This requires testing in rodent models of type 1 and type 2 diabetes and in rodents with defective CRR and optimisation of the dose to amplify CRR without worsening fasting/fed hyperglycemia. Development of brain permeable allosteric activators of AMPK could be useful for prevention of hypoglycemia in diabetes.

## METHODS

### Cell culture

Immortalised GT1-7 mouse hypothalamic cells were a kind gift from Pamela Mellon, Salk Institute, San Diego, California, USA. See supplementary methods for culture details. For experimentation, cells were incubated in serum-free DMEM containing physiologic brain glucose levels (2.5 mmol/L glucose). Cell lines were confirmed as mycoplasma free using a commercial kit (MycoAlert, Lonza, Slough, UK).

### Immunoblotting

Cells were grown to 60-70% confluence in 60 mm round petri dishes. Following treatment, cells lysates were collected for protein quantification using the Bradford method (27), separated using SDS-PAGE and transferred to nitrocellulose membrane. Total and phosphorylated protein was detected and semi-quantified using infrared fluorescence on the Licor Odyssey scanner. See supplemental methods for antibody details.

### Analysis of cellular metabolism

Mitochondrial oxygen consumption rates (OCR) were measured using the XF^e^96 Agilent Seahorse Extracellular Flux analyzer, as previously described (28). See supplemental methods for additional details.

### Determination of ATP concentrations

Cells were grown in 96-well plates overnight and intracellular ATP concentrations were measured using the ATPlite two-step assay (PerkinElmer, UK) as per manufacturer’s instructions and as previously described (28).

### Animals

Male Sprague-Dawley rats (250-350 g, Charles River Laboratories, Margate, Kent, UK) were maintained on a 12-hour light cycle, temperature 22-23 °C, 55% humidity and provided with food (Lab Diet; catalogue number 5LF2) and water *ad libitum*.

### Glucose tolerance tests

Male Sprague Dawley rats were fasted overnight. For studies using SBI-0206965 (3 mg kg^-1^; i.p.) and hexamethonium (50 mg kg^-1^; i.p.), either drug was delivered 30 minutes before glucose (2 g kg^-1^; i.p.) +/-R481 (5-20 mg kg^-1^; i.p); R419 (20 mg kg^-1^) or vehicle (0.5 % HPMC + 0.1 % TWEEN-80) in a single injection. Blood glucose measured at 0, 15, 30, 60 and 120 minutes from a tail vein prick by handheld glucometer (AccuCheck, Roche).

### Hyperinsulinemic clamp studies

Male Sprague-Dawley rats with pre-implanted jugular vein and carotid artery catheters were purchased from Charles River (Margate, UK). Rats were fasted overnight (16 hours). R481 (20 mg kg^-1^; i.p.; Rigel Pharmaceuticals) or vehicle (0.5% HPMC, 0.1% TWEEN-80 prepared in distilled sterile H_2_O) was administered one hour prior to the hyperinsulinemic-euglycemic or hypoglycemic clamp. Blood glucose was measured every 5-10 min and larger blood samples for hormone analysis were collected every 30 min.

### Hormone and metabolite analysis

Plasma glucagon and C-peptide were measured using the Mercodia Glucagon and C-peptide ELISA kits (Uppsala, Sweden). Plasma epinephrine was measured using the Demeditec Adrenaline ELISA (Kiel, Germany).

### Statistical analysis

A one-sample t-test was used to determine significant changes in phosphorylation or total protein expression relative to control in immunoblotting experiments. Plasma glucose levels, glucose infusion rates and plasma analytes were analysed using a two-way ANOVA with repeated measures. Peak hormone levels were analysed using an unpaired t-test. Analysis was performed using the GraphPad Prism software (Prism 8, GraphPad Software, La Jolla, CA, USA). Results are expressed as mean ± SEM, with *p < 0.05* considered statistically significant.

### Study approval

All procedures were approved by the University of Exeter Animal Welfare and Ethical Review body and were performed in accordance to the Animals Scientific Procedures Act (1986).

## AUTHOR CONTRIBUTIONS

A.M.C, Y.M., J.M.V.W., P.G.W.P, K.P, and C.B. researched data. A.M.C., J.M.V.W., S.S., K.L.J.E., and C.B., contributed to study design. All authors contributed to writing the manuscript and approved the final version. C.B. conceived the study, had access to all the data collected at the University of Exeter and takes responsibility for the accuracy and integrity of the data.

## ACKNOWLEDGEMENTS

This study was funded by: a JDRF Innovative grant (1-INO-2016-214-A-N) to C.B. and K.L.J.E; a Diabetes UK RD Lawrence Fellowship to C.B. (13/0004647); a Society for Endocrinology early career grant to C.B. and a British Society for Neuroendocrinology practical skills grant to C.B. A.M.C. is funded by a University of Exeter Medical School PhD studentship. We wish to thank Jennifer Gallagher, Dr Alison McNeilly, Prof Rory McCrimmon and Gary Park. We also wish to thank Prof René Remie for surgical refinements.

---

**Supplemental Figure 1.**
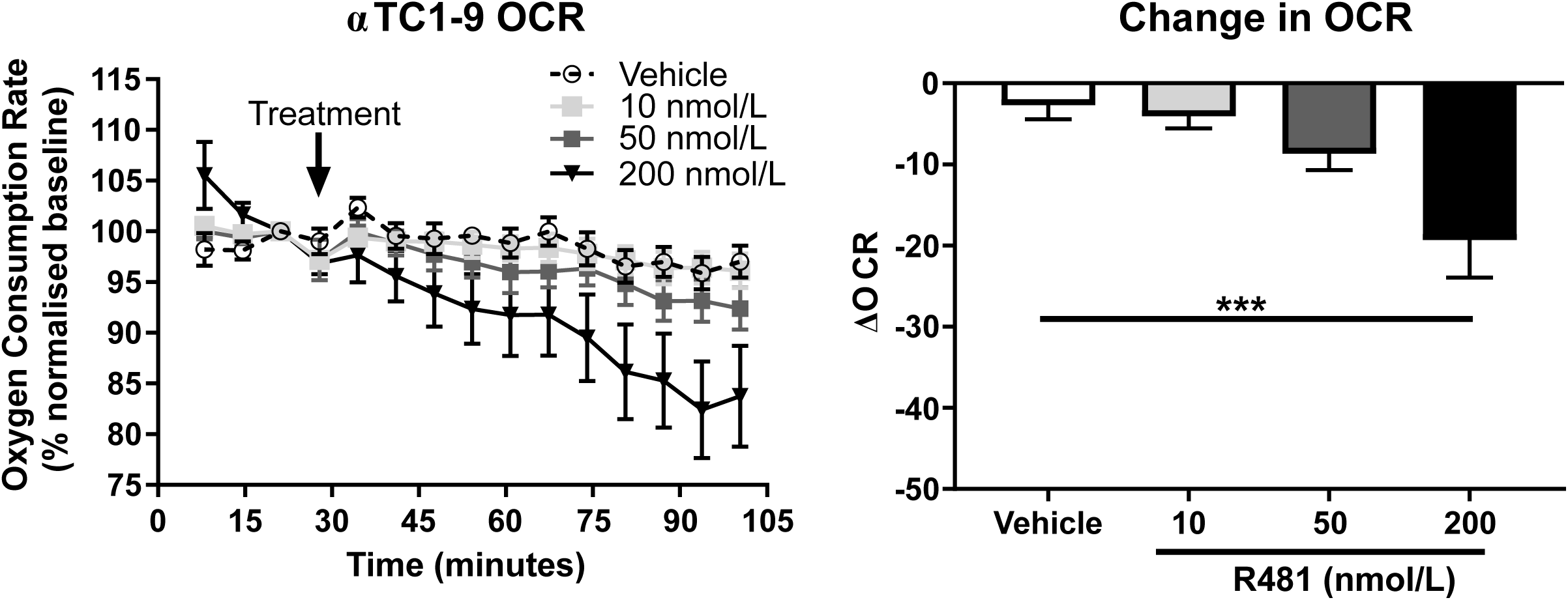
R481 reduces mitochondrial oxygen consumption in alpha cell line αTC1-9. **A.** Extracellular flux analysis measured by Seahorse analyzer (n=6-10). **B**. Change in Oxygen Consumption Rate (OCR) approximately 80 minutes post injection (denoted by arrow; n=6-10; ***P<0.005). Mean and SEM.

**Supplementary Figure 2.**
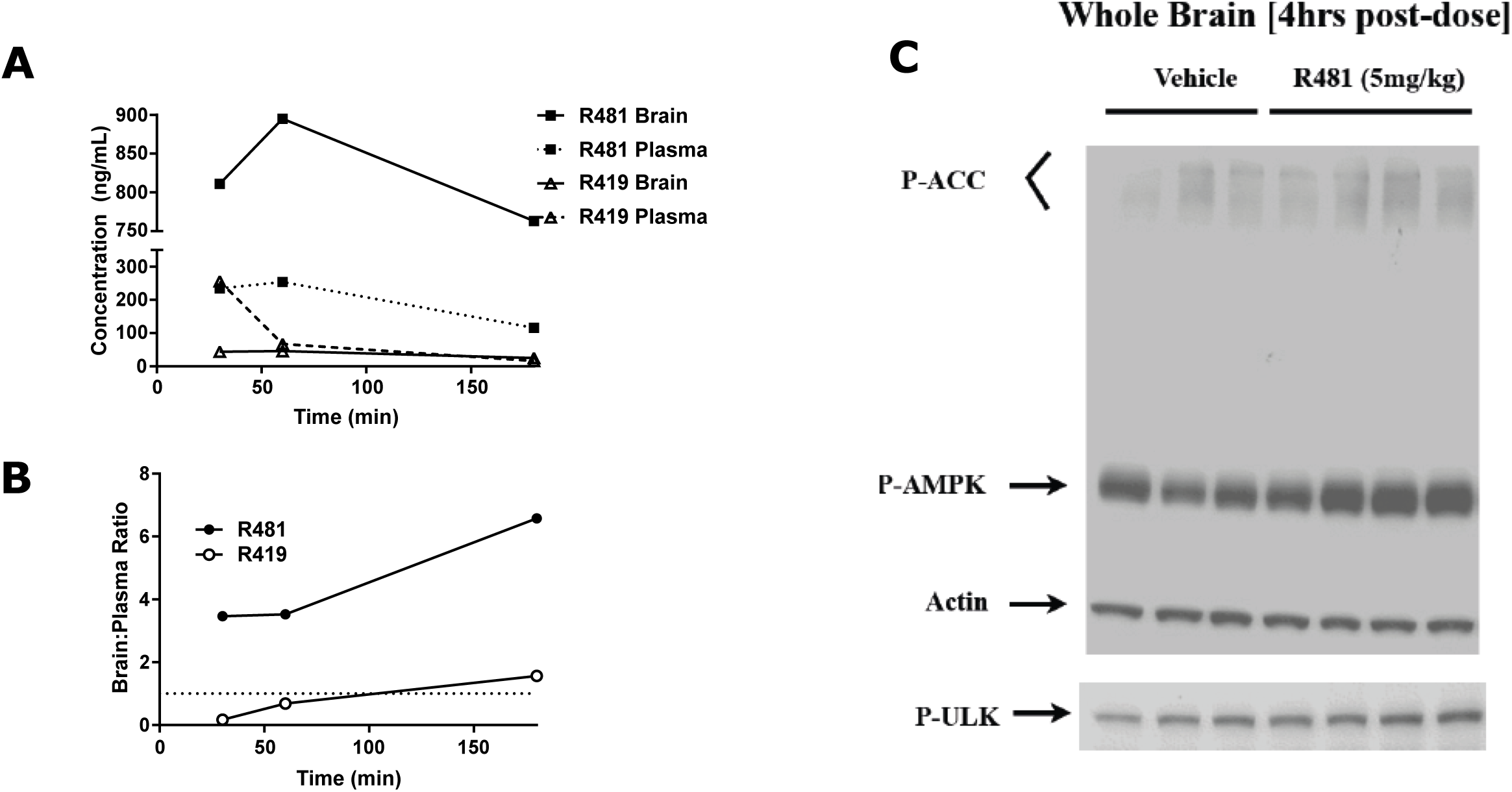
R481 accumulates in brain and activates AMPK in Balb/c mice. **A.** Brain and plasma levels of R481 and R419 measured from Balb/c mice following dosing (1 mg/Kg; i.v. bolus) **B.** Brain:plasma ratios of R48y and R4y9L **C.** Representative Western blots of pTy7I AMPK5 pS79 ACC and pS555 ULK1 4 hours after oral dosing in mice (R481; 5 mg/kg; p.o.), Data provided by Rigel Pharmaceuticals, Inc.

**Supplemental Figure 3.**
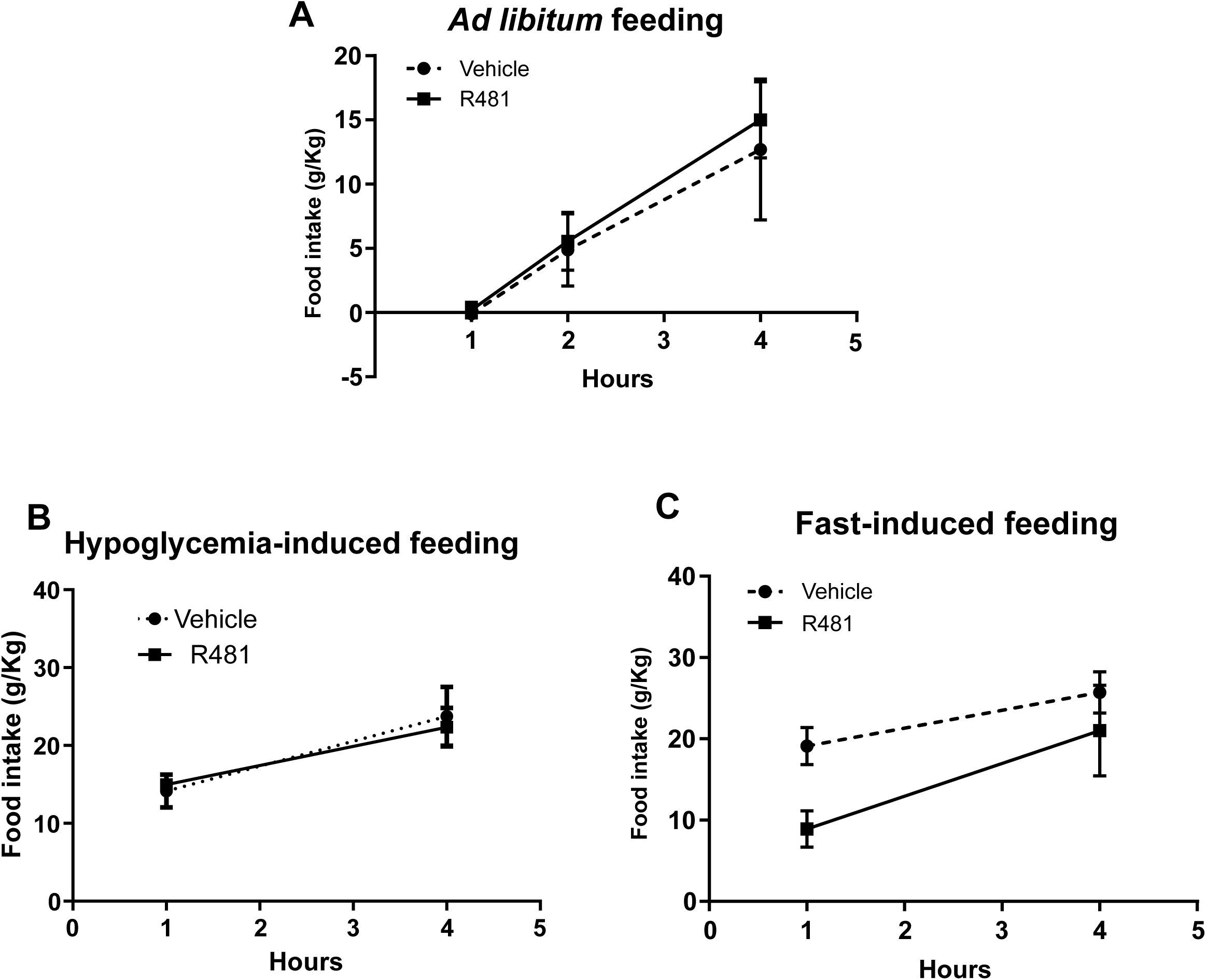
R481 had no effect on feeding ad libitum or in response to fasting or insulin treatment. **A.** R481 (20 mg kg^-1^:i.p.) was administered at the start of the dark-phase and food intake measured 1-4 hours following injection (Vehicle n=4; R48M n=4) **B.** R481 (5 mg kg^-1^:orally) was administered one hour prior to insulin injection (0.75 U kg^-14^ i.p.), food re-introduced after 6E minutes and intake measured M and 4 hours later (Vehicle n=5; R481 n=5) **C.** Rats were administered R481 (20 mg kg^-1^;i.p.) following M6 hour fast and food introduced immediately following injection and measured 1 and 4 hours later (Vehicle n=9; R481 n=9); TwoDway ANOVA with repeated measures. Data presented as mean ± SEM.

