## Supplemental Materials and Methods for "Brain permeable AMPK activator R481 raises glycemia by autonomic nervous system activation and amplifies the counterregulatory response to hypoglycemia in rats"

**Cell culture**

GT1-7 cells were cultured in media (Dulbecco’s modified eagles media [DMEM]; catalogue number D5671, Sigma, UK) containing 25 mmol/L glucose, supplemented with: 10% fetal bovine serum (FBS; Gibco, catalogue number 10270-106) and penicillin/streptomycin (Gibco; 100 U/ml; 100 μg/ml, respectively), as previously described (25). αTC1-9 cells were cultured in DMEM (catalogue number 31885, ThermoFisher, UK), containing 5.5 mmol/L glucose, supplemented with: 10% FBS (catalogue number EU-000, Sera Labs;UK); penicillin/streptomycin (Gibco; 100 U/ml; 100 μg/ml, respectively); HEPES (15 mmol/L; catalogue # H0877; Sigma, UK), Mixed ssential Amino Acids (catalogue # 11130051; ThermoFisher, UK); 0.02% (w/v) Factor V Fatty Acid Free Bovine Serum Albumin (catalogue number 10775835001, Sigma, UK ). Cells were maintained at 37**°**C, 5% CO_2_ and atmospheric O_2_ in a humidified incubator and confirmed as mycoplasma free using a commercial kit (MycoAlert, Lonza, Slough, UK).

**Immunoblotting**

Primary antibodies used were: pThr172 AMPK (1:1,000; catalogue #2535), pSer79 acetyl CoA carboxylase (ACC; 1:1000; catalogue #3661) from Cell Signaling Technologies, total ACC (1:1,000; catalogue #05-1098) from Merck Millipore and β-actin (1:10,000; catalogue #ΝΒ600-501) from Biotechne.

**Analysis of cellular metabolism**

GT1-7 cells were seeded on to XFe 96 well plates at 30,000 cells per well the day prior to assay, in wells pre-coated with poly-L-lysine (4 µg/mL) to enhanced cellular adhesion to surfaces. Cells were maintained in DMEM containing 7.5 mmol/L glucose as above. The day of study, cells were incubated in serum-free DMEM for 2 hours prior to incubation in low buffered media containing 2.5 mmol/L glucose (pH 7.4 at 37 degrees) for one hour in a non-CO_2_ incubator. R481 was injected after the 4^th^ mix-measure cycle and respiration monitored for approximately 80 minutes. Cellular material was collected in NaOH (50 mmol/L) for protein quantification using the method of Bradford. OCR values (pmol/min) were normalised to protein content then to baseline levels on the 4^th^ mix-measure cycle, prior to R481/vehicle injection. αTC1-9 cells were seeded at 70,000 cells per well and incubated as above, with the exception that glucose concentrations during the assay were maintained at 5.5 mmol/L.

**Hyperinsulinemic clamp studies**

Male Sprague-Dawley rats were surgically implanted with carotid artery and jugular vein catheters at Charles River (Margate UK). Catheters were exteriorised using a dual-channel vascular access button (Instech, USA) and covered using a lightweight aluminium cap, allowing, where possible, social housing following surgery. Catheter patency was maintained by flushing the catheters every 3-4 days with heparinized glucose catheter lock solution. During euglycemic clamps (120 minutes in total), animals received a fixed continuous insulin infusion of 20 mU kg^-1^ min^-1^ and a variable dextrose (50% w/v) to maintain glycaemia at approximately 5.5 mmol/L. To induce hypoglycemia, rats received a bolus insulin infusion of 80 mU kg^-1^ min^-1^ for 10 minutes, followed by a maintenance dose of 20 mU kg^-1^ min^-1^ for the remainder of the clamp (90 minutes in total). A variable 20% (w/v) dextrose infusion was used to maintain blood glucose levels at approximately 2.8 mmol/L at nadir.
